# Shifting flood regimes alter iron–sulfur metabolism and greenhouse gas associations

**DOI:** 10.1101/2025.03.03.641170

**Authors:** Colin G. Finlay, Ariane L. Peralta

## Abstract

Coastal wetlands store carbon, but compound stressors, including saltwater intrusion, sea level rise, and precipitation extremes, threaten this benefit by altering microbial communities and influencing greenhouse gas emissions. Mediating soil gas exchange, nutrient and carbon availability, and soil moisture, wetland vegetation interacts with these compound stressors. However, the exclusion of plant-driven redox shifts from many mesocosm studies limits our understanding of their role in shaping microbial responses to hydrologic conditions. This study employed a soil mesocosm approach to investigate the impact of hydrology (wet, dry, and interim) and plant presence on microbial communities and greenhouse gas fluxes in coastal freshwater wetland soils with varying past hydrologic regimes (i.e., soil history) and salinity stress. We used shotgun metagenomic sequencing to characterize the functional potential of soil microbes, measured *in situ* greenhouse gas fluxes, and characterized soil physicochemistry. Results showed that contemporary hydrology and soil history significantly impacted microbial gene composition related to sulfate and iron reduction. The compositions of genes for sulfate and iron reduction were correlated, and dissimilatory sulfate reduction genes influenced methane emissions. Findings highlight the roles of historical hydrology, potential saltwater exposure, and soil iron in shaping microbial responses to future changes in soil moisture, plant cover, and salinity. While plants did not significantly influence sulfur or iron metabolism, plant presence did impact green-house gas fluxes. We found a strong relationship between sulfate reduction and methanogenesis, which complements previous studies that have shown enhanced methanogenesis with seawater amendment. These results indicate that flooding without salinity is sufficient for coupled sulfate reduction and methanogenesis, provided that a legacy of saltwater intrusion has altered soil sulfate concentrations and sulfate-reducing microbial communities. Understanding microbial community metabolism in coastal wetlands is crucial for predicting their role in carbon sequestration and greenhouse gas emissions under future climate scenarios, developing effective management strategies to mitigate climate change impacts, and preserving these vital wetland ecosystems.

## 1. Introduction

Coastal freshwater wetlands are major sinks for atmospheric carbon (C) and reactive nitrogen (N) (Kayranli et al., 2010; Jordan et al., 2011; Nahlik and Fennessy, 2016). But these ecosystem services are threatened by saltwater intrusion and sea level rise (SWISLR) (Hopkinson et al., 2012; Wilson et al., 2018; Neubauer et al., 2019; Widney et al., 2019), changing precipitation patterns (Chu et al. 2019; Wang et al. 2024; Liu et al. 2025), and loss of wetland vegetation (Jordan et al., 2011; Nahlik and Fennessy, 2016). SWISLR can impose flooded conditions and the influx of terminal electron acceptors (TEA) such as sulfate (SO_4_^2−^) (Hopfensperger et al., 2014; Herbert et al., 2015). Flooding alters the TEA pool, by decreasing soil oxygen (O_2_) levels (Kozlowski, 1984), and saltwater intrusion from marine sources introduces SO_4_^2−^ to the ecosystem (Ardón et al., 2013; Schoepfer et al., 2014; Herbert et al., 2015). The combined impacts of flooding and SO_4_^2−^ addition on wetland ecosystem function are difficult to predict, in part because varying environmental characteristics, especially soil ferrous iron (Fe(II)), could buffer the effects of SO_4_^2−^ intrusion (Schoepfer et al., 2014; Johnson et al., 2019; Li et al., 2024). With climate change predicted to increase rates of sea level rise and increase the frequency of storm surges from more intense tropical storms, we need to better understand how the combined environmental stressors, saltwater intrusion and flooding, influence coastal wetlands (IPCC, 2023).

To harvest energy through respiration, microorganisms oxidize organic or inorganic substances and transfer the electrons through electron transport chains to ultimately reduce a TEA (reviewed in Burgin et al., 2011). O_2_ is used as a TEA in aerobic respiration (reviewed in Glass et al., 2022); in the absence of O_2_, a variety of other TEAs may be used in anaerobic respiration, including nitrate (NO_3_ ^−^), ferric iron (Fe(III)), SO_4_^2−^, and carbon dioxide (CO_2_) (Wang et al., 2017) (Figure S1). In common with these respiratory (i.e., dissimilatory) processes, assimilatory processes also involve oxidation-reduction (redox) reactions (Burgin et al., 2011). The availability of specific reductants (e.g., organic matter) and specific oxidants (e.g., TEAs like O_2_) modulates both dissimilatory and assimilatory processes (Burgin et al., 2011). The soil redox potential, which is the overall propensity of a soil environment to reduce or oxidize, integrates relative levels of reductants and oxidants within the soil (Burgin et al., 2011). In this study, we also refer to the overall oxidation-reduction propensity of the environment as “redox status”.

Soil redox potential influences microbial community structure and function (DeAngelis et al., 2010; Peralta et al., 2014). As soil microbes are major drivers of biogeochemical cycles, soil redox potential affects microbial metabolic capacity, ultimately influencing transformations of C, N, sulfur (S), and Fe at the ecosystem level (Falkowski et al., 2008; Schlesinger et al., 2011; Wang et al., 2017; Crowther et al., 2019). While the prediction of microbial metabolic diversity is inferred by taxonomic approaches, characterizing microbial metabolic diversity by examining genetic potential (e.g., shotgun metagenomics) provides a more direct evaluation of functional potential (Quince et al., 2017; Toole et al., 2021). To further improve the prediction of microbial functions, it is critical to understand how changing environmental conditions directly affect soil redox conditions.

Dynamic wetland hydrology modifies soil redox status (Peralta et al., 2013, 2014). Flooding of soils fills pore space, thereby restricting access to atmospheric O_2_ (Pezeshki and DeLaune, 2012). This creates suboxic and anoxic conditions, requiring alternative TEAs to be used in anaerobic respiration (Patrick Jr. and Jug-sujinda, 1992; RoyChowdhury et al., 2018) (Figure S1). Based on the hydrologic regime of a wetland, soils therein may experience fluctuating wet-dry conditions, as well as prolonged inundation or dry down (DeAngelis et al., 2010; Peralta et al., 2014; RoyChowdhury et al., 2018).

In addition to dynamic hydrology, human activities modify biogeochemical cycles. For example, watersheds accumulate N via human activities, and this N may be loaded in coastal wetlands via hydrologic flows and atmospheric deposition (Conley et al., 2009; Chilton et al., 2021). The incoming anthropogenic N may be in the form of NO_3_ ^−^ or converted to NO_3_ ^−^ by internal microbial processing (e.g., nitrification, NH_4_^+^ → NO_3_ ^−^) (reviewed in Kuypers et al., 2018). Since NO_3_ ^−^ is an efficient TEA, it is used in facultative anaerobic respiration, which is a common metabolism observed in fluctuating oxic/anoxic conditions (Wallenstein et al., 2006). In addition, saltwater intrusion events can increase soil and porewater SO_4_^2−^ concentrations (Herbert et al., 2018). Then, SO_4_^2−^-reducing microorganisms (SRM) produce sulfide which binds with reduced Fe to form iron sulfide (FeS) (Schoepfer et al., 2014). This buffering of seawater SO_4_^2−^ is, therefore, dependent on the activity of SRM and Fe-reducing microorganisms (FeRM), as well as the existing soil Fe concentrations (Schoepfer et al., 2014).

While human activities influence biogeochemical cycles, cross-kingdom interactions between plants and microorganisms also have important consequences for soil redox activity. Plants alter soil redox potential by delivering O_2_ to the rhizosphere and transporting CO_2_ and methane (CH_4_) from the rhizosphere to the atmosphere (Colmer, 2003). Plant root O_2_ release can buffer the effects of flooding on soil aeration (Kozlowski, 1984; Cook and Knight, 2003; Koop-Jakobsen and Wenzhöfer, 2015). While hydrologic status can be variable over time, plants impose a more constant influence on soil redox conditions (Koop-Jakobsen and Wenzhöfer, 2015).

The biogeochemical cycles of C, N, Fe, and S are not isolated, but instead, closely linked. For example, NO_3_ ^−^ reduction can be coupled to sulfide oxidation (Burgin and Hamilton, 2008). The oxidation of organic C is coupled to reductions of NO_3_ ^−^, Fe(III), and SO_4_^2−^ (Burgin et al., 2011). The microbial N cycle forms an interconnected network, with many reactions dependent on the availability of C, Fe, and S (Kuypers et al., 2018). Therefore, to understand the cycling of any one element, it is essential to consider the relevant microbe-microbe, plant-microbe, and microbe-environment interactions that influence the nutrient cycling network. These microbial nutrient cycling networks provide crucial context for understanding specific environmental phenomena, such as SWISLR and Fe buffering of ecosystem sulfidization. Past work on saltwater intrusion and Fe-S buffering was conducted at the Timberlake Observatory for Wetland Restoration (TOWeR) (Schoepfer et al., 2014). The TOWeR site is prone to seasonal, drought-induced saltwater intrusion, which introduces SO_4_^2−^ to Fe-rich soils (Ardón et al., 2013; Lamers et al., 2013; Schoepfer et al., 2014). Both SO_4_^2−^ and Fe can be microbially reduced to sulfide and Fe(II), respectively (Weber et al., 2006; Lamers et al., 2013; Schoepfer et al., 2014). The sulfide may be toxic to salinity-na**ï**ve ecosystems, but can readily bind Fe(II) to create FeS, which is non-toxic and relatively non-bioavailable (Lamers et al., 2013; Schoepfer et al., 2014). For example, Schoepfer and colleagues (2014) found that the pool of reduced Fe in TOWeR soils is sufficient to sequester sulfide from seasonal saltwater intrusion into FeS, but as SWISLR accelerates soil sulfidization, toxic sulfide levels may accumulate in the future.

How future sulfidization (due to SWISLR) will modify wetland microbial community structure and function is unknown. Specifically, it is unclear how wetland microbes that participate in S cycling will respond to flooding, drought, and changes in plant cover. This study used a manipulative mesocosm approach to address the questions (1) how do soil Fe and S concentrations interact with altered redox states (hydrology and plants) and microbial community composition of SO_4_^2−^ reduction genes? and (2) how do these Fe-S metabolic associations relate to greenhouse gas (GHG) emissions, mediated by C and N metabolisms? We hypothesize that the most reducing conditions (i.e., prolonged flooding, no plants) modify different anaerobic metabolisms in similar ways and predict that (i) in oxidizing conditions (dry and/or plant presence) the composition of microbes that could participate in SO_4_^2−^ reduction and Fe reduction will not be linked/coupled. Coupled here means a significant correlation (i.e., Mantel test) between Bray-Curtis distance matrices of SO_4_^2−^ reduction and Fe reduction genes. We also pre-dict that (ii) in reducing conditions (wet and/or plant absence) coupling between the composition of microbes that could participation in SO_4_^2−^ reduction and Fe reduction will be observed, and these processes will contribute to CO_2_ production (fit with CO_2_ fluxes using envfit) while competing with methanogenesis (observed as negative relationship with methanogenic functional genes and CH_4_ fluxes).

## 2. Methods

### 2.1. Sampling and mesocosm experimental design

On April 1, 2016, soil samples were collected from TOWeR located in the Albemarle Peninsula in Tyrell County, North Carolina, USA (35°54’22” N, 76°09’25” W) (Ardón et al., 2010; Morse et al., 2012). The site is connected to the Albemarle Sound via the Little Alligator River, with potential for saltwater intrusion (Ardón et al., 2013). Across the sampling area, the position of the water table creates a hydrologic gradient, with sites categorized as dry (upland), wet (saturated), and interim (transition dry/wet) (Hopfensperger et al., 2014) (Figure 1).

**Figure 1.**
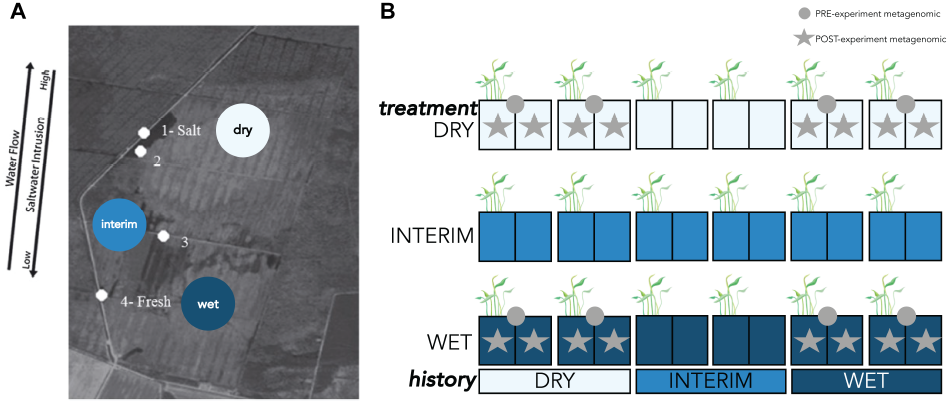
Field site sampling and experimental design of the wetland mesocosm experiment. Wetland mesocosms sourced from the Timberlake Observatory for Wetland Restoration field site (image modified from Schoepfer et al. (2014)), with hydrologic histories for the present study (dry, interim, wet) labeled with circles (A). Numbered white circles on the map indicate the sampling locations from Schoepfer et al. (2014), and the determination of salt and freshwater conditions during sampling in 2012. The schematic on the right shows each replicate of the mesocosm experiment, with grey circles representing mesocosms sampled for pre-experiment metagenomics (n = 8), and grey stars representing mesocosms sampled for post-experiment metagenomics (n = 16).

Soil blocks collected from the three sites within TOWeR (dry, wet, interim) were used to set up a mesocosm experiment. Soil history was categorized according to water table levels at the time of collection (dry = 20 cm, interim = 10 cm, and wet = 0 cm below surface level). Six intact soil blocks (25 cm x 25 cm x 20 cm) from each of the three hydrologic histories were contained in dark plastic containers of the same dimensions. Soil mesocosms were maintained for the duration of the experiment at the East Carolina University (ECU) West Research Campus (Greenville, NC) under a covered hoop house with a shade cloth to prevent precipitation while allowing 60% light penetration. Each block was divided in half by a root screen (20 µm stainless steel mesh). Plants were allowed to grow on one side of the screen and removed (above- and belowground) on the other side of the screen to create “plant” and “no plant” treatments. Polyvinyl chloride (PVC) collars were installed in each mesocosm as a base for GHG sampling.

Hydrologic treatments were started two weeks after the installation of PVC collars and the initiation of plant and no plant treatments. To manipulate hydrology, vinyl tubing connected to a 1-L water bottle was inserted on each side of the mesocosm, to a depth of three cm from the bottom of the mesocosm container. Water levels inside the mesocosm were maintained by filling the 1-L bottle to the desired height using rainwater collected from a cistern. Mesocosms exposed to wet conditions were flooded and maintained at maximum container height (18 cm). Mesocosms exposed to dry conditions were maintained at a five cm water level. Interim hydrologic treatment fluctuated between flooded and dry conditions every two weeks, starting with a wet treatment and ending with a dry treatment. Experimental treatment lasted eight weeks, with 36 mesocosms evenly representing three levels of soil history (dry, wet, interim), two levels of plant treatment (plant, no plant), and three levels of hydrologic treatment (dry, wet, interim) in a factorial experimental design (Figure 1).

### 2.2. Measuring greenhouse gases

We investigated the impact of hydrology and vegetation on GHG production, measuring GHG fluxes starting on June 13, 2016, two weeks after establishing hydrologic treatments, and continuing biweekly until August 11, 2016, for a total of five sampling events. GHG collection chambers, made from clear acrylic tubing, were 20 cm high with a diameter of 8.25 cm. Using clear chambers allowed us to capture total GHG fluxes, not just soil respiration due to microbial activity. Chambers were sealed with silicon and a cap fitted with a 33 mm septum as a gas sampling port. During sampling, the chambers were placed on pre-installed PVC collars and sealed with tape to prevent diffusion (Hoffmann et al., 2018). Gas samples were collected 30, 60, 120, and 180 minutes after chamber attachment. Ambient (T_0_) samples were not collected before chamber closure but were later collected from the mesocosm incubation field site (ECU West Research Campus) as part of a 16-month GHG sampling effort, from August 2021 to December 2022. Monthly average ambient GHG concentrations were calculated and used as the T_0_ concentration for the mesocosm flux time points in the corresponding month. All GHG samples were collected using a 20 mL syringe with a needle, mixing the headspace gas three times before collecting a 20 mL sample, then splitting the sample between a pair of three mL glass Exetainer vials (Lampeter, Wales, UK) with screw tops and dual-layer septa. Samples were stored upside down at room temperature in a dark location and analyzed within 96 hours.

We analyzed GHG concentrations with a Shimadzu gas chromatograph (GC-2014), equipped with an electron capture detector for nitrous oxide (N_2_O) and a flame ionization detector with a methanizer for CH_4_ and CO_2_. Calibration standards have been described previously (Bledsoe et al., 2025). Samples exceeding the calibration curve were diluted and reanalyzed. GHG fluxes were calculated using the gasfluxes package implemented in the R Statistical Environment and RStudio (RStudio 2025.05.0+496, Rv4.5.0) (Fuss and Hueppi, 2024; Posit Team, 2025; R Core Team, 2025).

### 2.3. Amplicon and shotgun metagenomic sequencing

Based on amplicon sequencing, we identified a subset of samples representing the most distinct microbial communities for shotgun metagenomic sequencing (Peralta et al., 2020; Bledsoe et al., 2025). Briefly, we chose a set of baseline (before hydrologic/plant treatment) and post-flooding/drying treatment samples. Baseline samples were sourced from dry (water level approximately −20 cm, n = 4) and wet (water level approximately 0 cm, n = 4) hydrologic conditions, referred to as soil history. At the end of the eight-week experiment, we selected a subset of samples from the hydrologic treatments (prolonged drying or wetting only) and plant treatments (presence or absence of vegetation) (n = 16). After genomic DNA extraction using the Qiagen DNeasy PowerMax Soil Kit, samples were sent to the U.S. Department of Energy (DOE) Joint Genome Institute (JGI) for sequencing and analyses (GOLD study ID Gs0142547 and NCBI BioProject accession number PRJNA641216). Bin methods in the IMG pipeline (MetaBAT version 2.12.1, CheckM v1.0.12, GTDB database release 86, GTDB-tk version v0.2.2) were used to curate 14 medium-to high-quality bins from the metagenomes. Medium-quality bins have at least 50% completion and less than 10% contamination. High-quality bins have greater than 90% completion, less than 5% contamination, the presence of 23S, 16S, and 5S rRNA genes, and at least 18 tRNA genes (Bowers et al., 2017). Further description of sample processing and metagenomic sequencing can be found in Peralta et al. (2020).

From the metagenomes, we curated functional gene sets using IMG/M’s integrated Kyoto Encyclopedia of Genes and Genomes (KEGG) module list (Chen et al., 2017; Kanehisa and Goto, 2000). We focused on the following four functional gene sets associated with S cycling: Assimilatory Sulfate Reduction (M00176), Dissimilatory Sulfate Reduction (M00596), Thiosulfate Oxidation by SOX Complex (M00595), and Sulfate-Sulfur Assimilation (M00616). We also analyzed KEGG module M00530: Dissimilatory Nitrate Reduction to Ammonia (DNRA). Previously, we examined metabolic gene modules related to biogenic GHG emissions, including Methanogen (M00617) (Bledsoe et al., 2025). For a comparison of functional genes impacting CH_4_ flux, we analyzed Methane Oxidation (M00174).

The IMG/M and publicly available databases in general have limited options for Fe gene analysis. From IMG/M’s integrated TIGRFAM tool, we analyzed two genes putatively involved in Fe reduction: decaheme c-type cytochrome, OmcA/MtrC family (TIGR03507), and decaheme-associated outer membrane protein, MtrB/PioB family (TIGR03509). These two genes were analyzed together as one “module”, analogous to collections of KEGG Orthologs (KO) within each KEGG Module.

For further Fe gene analysis, we analyzed all metagenomes using FeGenie, a hidden Markov model (HMM) tool designed for the identification of Fe genes and Fe gene neighborhoods (Garber et al., 2020). Complete nucleotide assemblies (FNA files) for each metagenome were downloaded from IMG/MER, then searched using FeGenie’s custom HMM scripts. Bitscores greater than the bitscore cuttoff were counted as positive “hits” for a given Fe gene. Counts of gene hits were totaled within each of FeGenie’s gene categories: (1) iron acquisition (a. iron transport, b. heme transport, c. heme oxygenase, d. siderophore synthesis, e. siderophore transport, f. siderophore transport potential); (2) iron gene regulation; (3) iron oxidation; (4) possible iron oxidation and possible iron reduction; (5) probable iron reduction; (6) iron reduction; (7) iron storage; (8) magnetosome formation. Counts within each FeGenie category were then converted to relative abundance, and relative abundance matrices were converted to Bray-Curtis distance matrices. Functional gene counts within each respective KEGG Module and TIGRfam “module” followed a similar process: conversion of counts to relative abundance (relative to total module counts), and conversion of relative abundance matrices to Bray-Curtis distance matrices.

### 2.4. Measuring integrated soil redox status

Soil redox status was measured using Indicator of Reduction in Soils (IRIS) tubes, following the methodology described in our previous study (Jenkinson and Franzmeier, 2006; Bledsoe et al., 2025). At the beginning of the experiment, two IRIS tubes were installed in each mesocosm: one on the plant side and one on the bare soil (“no plant”) side. The tubes were incubated for two weeks before removal and analysis. A non-coated PVC pipe was used to fill the hole left by the removed IRIS tube. Two weeks before the end of the experiment, the non-coated PVC pipe was replaced with a new IRIS tube to measure soil redox status at the end of the experiment.

The surface area of Fe(III) paint removed from the IRIS tubes was quanti-fied using ImageJ (Schneider et al., 2012). The percentage of paint removed was used to interpret redox status, with higher percentages indicating more reducing conditions. For further details, please refer to Bledsoe et al. (2025).

### 2.5. Measuring soil factors

Soil properties were determined from soils collected during the destructive sampling of mesocosms at the end of the experiment. A sample of air-dried soil from each mesocosm was sent to Waters Agricultural Laboratories, Inc. (Warsaw, NC) and analyzed for pH, phosphorus, potassium, magnesium, sulfur, manganese, iron, and humic matter, using standard Mehlich III methods (Mehlich, 1984).

### 2.6. Statistical analyses

All statistical analyses were performed in the R statistical environment (RStudio 2025.05.0+496, Rv4.5.0) (Posit Team, 2025; R Core Team, 2025). We found GHG and IRIS tube data to be non-Gaussian and to have extreme outliers. To address these extreme outliers and non-normal data distribution, we employed nonparametric statistical methods to assess differences in GHG fluxes and IRIS percent across hydrologic treatments. For differences in GHG fluxes and IRIS percent among the three hydrologic treatments (dry, interim, and wet), we used Kruskal-Wallis Rank Sum tests, grouped by sampling date to accommodate repeated measures. When the Kruskal-Wallis test indicated significant differences among treatment groups for a given sampling date (p ≤ 0.05), we conducted pair-wise comparisons using the Dwass, Steel, Critchlow, and Fligner (SDCFlig) test using the function pSDCFlig in the package NSM3 (Schneider et al., 2024). We visualized bacterial community responses to soil history (field conditions), hydrologic treatment (contemporary dry/wet treatments), and plant presence/absence using the ordination method, principal coordinates analysis (PCoA), of bacterial community composition based on Bray-Curtis dissimilarity. We also used PCoA to visualize the composition of functional gene categories, separating points by soil history, hydrologic treatment, and plant presence/absence. We used permutational multivariate analysis of variance (PERMANOVA) to measure the variation in bacterial community composition due to soil history, hydrologic treatment, and plant presence. Hypothesis testing using PERMANOVA was performed using the vegan::adonis function (Oksanen et al., 2022). Finally, soil parameters and GHG fluxes were compared against bacterial community and functional patterns (based on Bray-Curtis dissimilarity) using the vegan::envfit function (Oksanen et al., 2022). The GHG fluxes and soil parameters with envfit *p* ≤ 0.05 were plotted as vectors, scaled by their correlation, on PCoA plots of functional gene composition. Distance-based partial least squares regression (DBPLSR) was used to measure relationships between Bray-Curtis distance matrices and GHG fluxes.

Next, we conducted a series of Mantel tests to measure correlations between Bray-Curtis distance matrices. We assessed the potential for NO_3_ ^−^-driven SO_4_^2−^ production by testing the distance matrix correlations between the composition of NO_3_ ^−^ reduction pathways (denitrification and DNRA) and the composition of thiosulfate oxidation by the SOX complex. We conducted a Mantel test for each respective NO_3_ ^−^ reduction pathway (denitrification and DNRA) correlated to thiosulfate oxidation by the SOX complex. We ran a series of Mantel tests for pairs of SO_4_^2−^ reduction, Fe reduction, and methanogenesis distance matrices. To investigate correlations between SO_4_^2−^ reduction, Fe reduction, and methanogenesis under different hydrological conditions, the modules were subset by soil history (wet or dry) and hydrologic treatment (wet or dry). We evaluated the strength of Mantel r correlations between pairs of SO_4_^2−^ reduction, Fe reduction, and methanogenesis distance matrices within each hydrologic condition (e.g., wet soil history). We used a statistical cutoff of p ≤ 0.05.

## 3. Results

Using a wetland mesocosm approach, we tested the following hypotheses: (i) in oxidizing conditions (dry and/or plant presence) SO_4_^2−^ reduction and Fe reduction will not be linked/coupled, where coupled means a significant correlation between Bray-Curtis distance matrices of SO_4_^2−^ reduction and Fe reduction; and (ii) in re-ducing conditions (wet and/or plant absence) coupling between SO_4_^2−^ reduction and Fe reduction will be observed, and these processes will contribute to CO_2_ production (fit with CO_2_ fluxes using envfit) while competing with methanogenesis (observed as negative relationship with methanogenic functional genes and CH_4_ fluxes). A summary of differences in community functional gene composition is reported in Table 1. The eight-week hydrologic manipulation resulted in differences in soil redox status based on the percentage of paint removed from IRIS tubes (IRIS percent) during the week zero and week eight time points measured at the start and end of the mesocosm incubation. During week zero, which followed two weeks of initial plant and hydrologic treatment and IRIS tube incubation, both interim and wet treatments exhibited more reducing conditions than the dry treatment (SDCFlig p ≤ 0.05; Figure S2). At the end of the eight-week experiment, the wet treatment displayed reducing conditions, whereas the dry and interim treatments did not (Figure S2).

**Table 1.** Summary of differences in community functional gene composition.

| Gene module | Plant | History | Treatment |
| --- | --- | --- | --- |
| Sulfate-Sulfur Assimilation |  |  | *<sup>a</sup> |
| Assimilatory Sulfate Reduction |  |  | * |
| Dissimilatory Sulfate Reduction |  | * |  |
| Sulfate Reduction (combined assimilatory and dissimilatory) |  | * | * |
| Thiosulfate Oxidation by SOX Complex |  | * |  |
| Fe Genes (FeGenie) |  | * |  |
| Fe Reduction (FeGenie) |  | * |  |
| Fe Reduction (TIGRFAMS mtrBC) | * |  | * |
| Dissimilatory Nitrate Reduction to Ammonium |  |  |  |
| Methane Oxidation |  |  |  |

<sup>a</sup> Permutational multivariate analysis of variance (PERMANOVA) was used to compare treatment differences in key functional categories, and differences in composition ( $p \leq 0.05$ ) are marked with an asterisk. This table is based on the complete PERMANOVA statistics found in Table S1.

### 3.1. Greenhouse gas fluxes

Here, we highlight general trends in GHG fluxes. A more detailed analysis of the GHG data is presented in (Bledsoe et al., 2025). The GHG fluxes varied over five time points collected every two weeks during the eight-week experiment. Soil history influenced CH_4_ fluxes during the first time point, with greater CH_4_ fluxes in mesocosms from the wet soil history compared to both interim and dry histories, and greater CH_4_ fluxes in interim histories compared to dry histories (Figure 2; Figure S3). At the first time point, N_2_O fluxes were greater in mesocosms with a wet soil history compared to those with an interim history (Figure 2; Figure S3). Differences in GHG fluxes between hydrologic histories were not observed after the first time point. Throughout the experiment, hydrologic treatment affected CH_4_ and CO_2_ fluxes, and the presence or absence of plants separated CO_2_ fluxes (Figure 2; Figure S4). While the CH_4_ fluxes were generally greater in the wet treatment compared to the dry treatment, CH_4_ fluxes were similar across treatments at both the first and final time points (Figure 2; Figure S4). The CO_2_ fluxes were greater in dry mesocosms and no plant vs. plant mesocosms, and the N_2_O fluxes were near zero and similar across treatments for most of the eight-week experiment (Figure 2; Figure S4). The trend in CH_4_ fluxes over time remained similar throughout the eight-week experiment 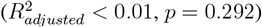, CO_2_ fluxes decreased slightly 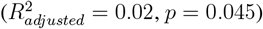, and N_2_O fluxes increased from slightly negative fluxes during the first two weeks, to near-zero fluxes during the final time points 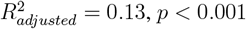; Figure 2).

**Figure 2.**
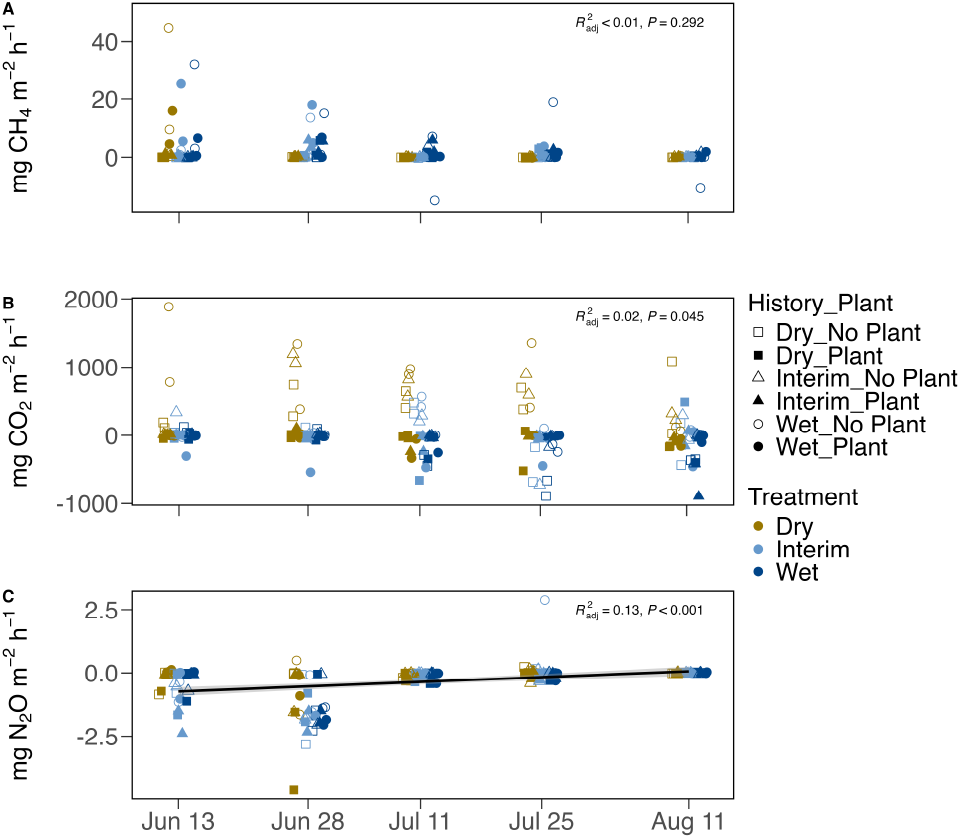
Greenhouse gas fluxes measured under different hydrologic conditions over time. Greenhouse gas (GHG) fluxes are in milligrams of greenhouse gas per square meter per hour. Individual fluxes are plotted as points and adjusted horizontally to avoid overlap. Color represents hydrologic treatment (brown = dry, light blue = interim, dark blue = wet). Shape represents soil history (square = dry history, triangle = interim history, circle = wet history), and closed shapes represent mesocosms with plants, while open shapes represent mesocosms without plants. A regression of GHG flux over time was performed, and a smoothed line of best fit is displayed (if *p* ≤ 0.05 and adjusted 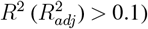) as a black line with 95% confidence intervals in grey. The adjusted R2adj and p-value for GHG flux as a function of time are displayed in the top right corner of each panel. An extreme outlier of 248 mg CH_4_ m^-2^ h^-1^, measured in a wet treatment with no plants on July 11^th^, was used in statistical calculations but not displayed in the plot to improve visualization of differences across treatments.

### 3.2. Sulfur functional genes

We measured how hydrology and plant presence influenced the community composition of genes involved in S cycling. Hydrologic treatment strongly influenced the composition of SO_4_^2−^-S assimilation (PERMANOVA, *F*_1,23_ = 3.917, *R*^2^ = 0.144, *p* = 0.008; Table 1; Table S1) and assimilatory SO_4_^2−^ reduction functional genes (PERMANOVA, *F*_1,23_ = 4.111, *R*^2^ = 0.156, *p* = 0.013; Table 1; Table S1). The response of SO_4_^2−^-S assimilation and assimilatory SO_4_^2−^ reduction gene composition correlated with redox status measured as the percent of paint removed from IRIS tubes (sulfate assimilation-IRIS envfit, *R*^2^ = 0.285, *p* = 0.032; Figure 3A; assimilatory sulfate reduction-IRIS envfit, *R*^2^ = 0.301, *p* = 0.027; Figure 3B). Soil history strongly influenced the compositions of genes involved in dissimilatory SO_4_^2−^ reduction (PERMANOVA, *F*_1,23_ = 10.397, *R*^2^ = 0.316, *p* = 0.0003; Table 1; Table S1) and thiosulfate oxidation by the SOX complex (PERMANOVA, *F*_1,23_ = 6.547, *R*^2^ = 0.243, *p* = 0.009; Table 1; Table S1). The combination of assimilatory and dissimilatory SO_4_^2−^ reduction differed in composition according to soil history and treatment (PERMANOVA, history: *F*_1,23_ = 2.830, *R*^2^ = 0.108, *p* = 0.030, treatment: *F*_1,23_ = 2.906, *R*^2^ = 0.111, *p* = 0.026; Table 1; Table S1). The community composition of SO_4_^2−^ reduction (assimilatory and dissimilatory) genes was significantly correlated with percent soil moisture (envfit, *R*^2^ = 0.298, *p* = 0.023; Figure 4C). The composition of dissimilatory sulfate reduction genes differed according to soil history (PER-MANOVA, history: *F*_1,23_ = 10.397, *R*^2^ = 0.316, *p* = 0.0003; Table 1; Table S1), and was related to CH_4_ flux (envfit, *R*^2^ = 0.401, *p* = 0.003; Figure 3C; Figure 5). Based on DBPLSR, the percent of variance in CH_4_ flux explained by dissimila-tory SO_4_^2−^ reduction decreased by 12% from week zero to week eight (adjusted 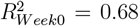, adjusted 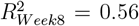, based on components 1 and 2; Figure 5; Table S2). Functional genes for CH_4_ oxidation and methanogenesis explained less variance in CH_4_ fluxes than dissimilatory SO4-reduction functional gene composition (Figure 5; Table S2). The relationship between CH_4_ fluxes and CH_4_ oxidation increased in strength during the experiment, and that of methanogenesis decreased from week zero to week eight (Figure 5; Table S2). The composition of methanogenesis functional genes was correlated with the compositions of genes involved in assimilatory SO_4_^2−^ reduction (Mantel *r* = 0.22, *p* = 0.01) and dissimilatory SO_4_^2−^ reduction (Mantel *r* = 0.26, *p* = 0.02; Figure 6). When we examined correlations between methanogenesis and SO_4_^2−^ reduction genes within hydrologic histories and treatments, we found assimilatory SO_4_^2−^ reduction and methanogenesis were most strongly correlated in dry treatment (Mantel *r* = 0.37, *p* = 0.01), while dissimilatory SO_4_^2−^ reduction and methanogenesis were most strongly correlated in wet treatment (Mantel *r* = 0.5, *p* = 0.01; Figure S5).

**Figure 3.**
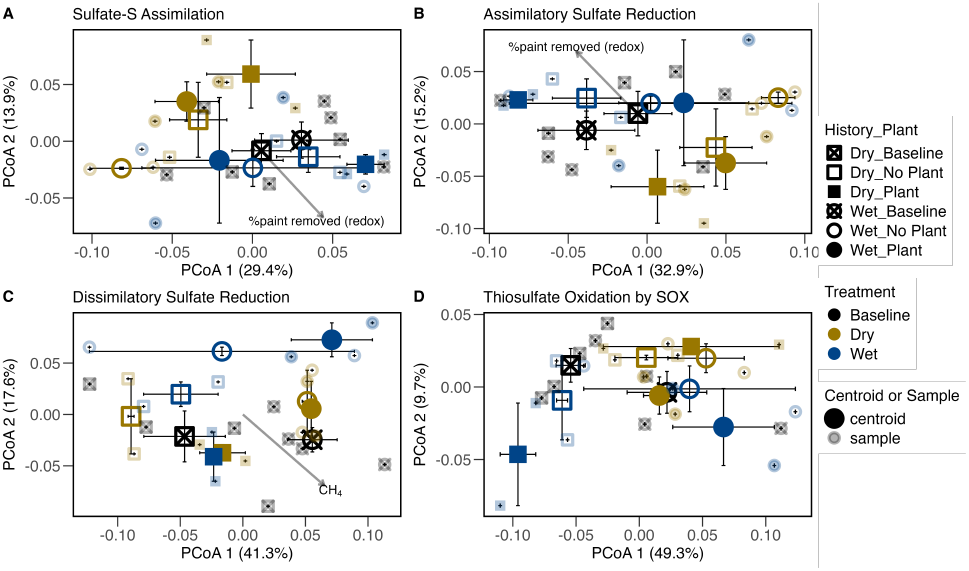
Ordinations of sulfur functional gene modules. Ordinations are based on principal coordinates analysis (PCoA), depicting community composition of the sulfur functional gene modules Sulfate-Sulfur Assimilation (M00616) (A), Assimilatory Sulfate Reduction (M00176) (B), Dissimilatory Sulfate Reduction (M00596) (C), and Thiosulfate Oxidation by SOX Complex (M00595) (D). The percent variance explained by each axis is listed in parentheses. Colors refer to hydrologic treatments, where black represents the baseline, brown represents dry conditions, and dark blue represents wet conditions. Shapes refer to the history of the sample, where squares represent dry history, and circles represent wet history. The shape fill represents plant treatment, where ‘x’ through symbol = baseline, open symbol = no plant, closed symbol = plant. Baseline samples were collected before the start of hydrologic and plant treatments, and treatment samples were collected after eight weeks. Vectors represent significant (*p* ≤ 0.05) correlation between greenhouse gas trends or soil redox status (as measured by the percent of paint removed from IRIS tubes) and functional gene composition, scaled by the magnitude of correlation (using envfit).

**Figure 4.**
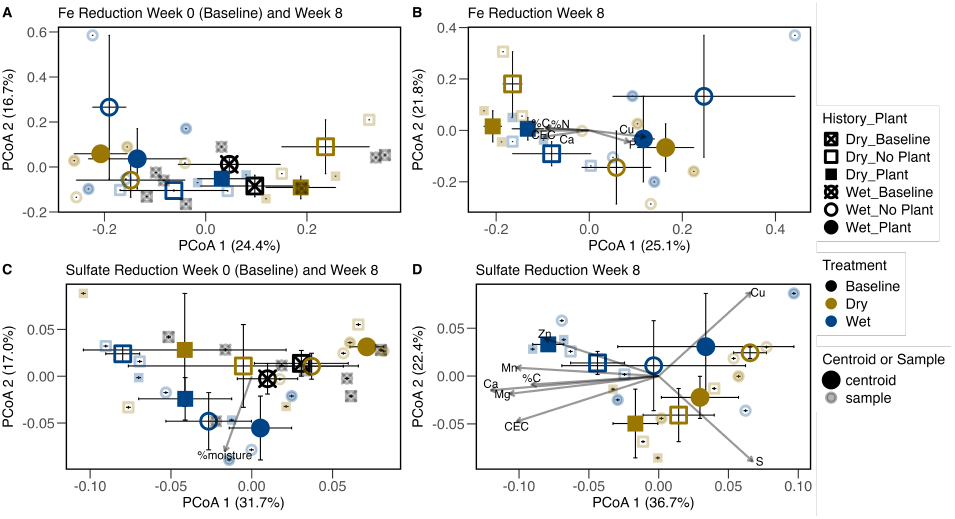
Ordinations of iron reduction and sulfur reduction genes. Ordinations are based on principal coordinates analysis (PCoA), depicting community composition of the functional gene modules: Fe reduction genes identified by FeGenie at Week 0 and Week 8 of the 8-week experiment (A), Fe reduction genes identified by FeGenie at end of 8-week experiment (B), sulfate reduction genes (both assimilatory and dissimilatory) at Week 0 and Week 8 of 8-week experiment (C), and sulfate reduction genes (both assimilatory and dissimilatory) at end of 8-week experiment (D). The percent variance explained by each axis is listed in parentheses. Colors refer to hydrologic treatments, where black represents the baseline, brown represents dry conditions, and dark blue represents wet conditions. Shapes refer to the history of the sample, where squares represent dry history and circles represent wet history. The shape fill represents plant treatment, where ‘x’ through symbol = baseline, open symbol = no plant, closed symbol = plant. Baseline samples were collected before the start of hydrologic and plant treatments, and treatment samples were collected after eight weeks. Vectors represent significant (*p* ≤ 0.05) correlation between greenhouse gas trends or soil physicochemical parameters, and functional gene composition, scaled by the magnitude of correlation (using envfit).

**Figure 5.**
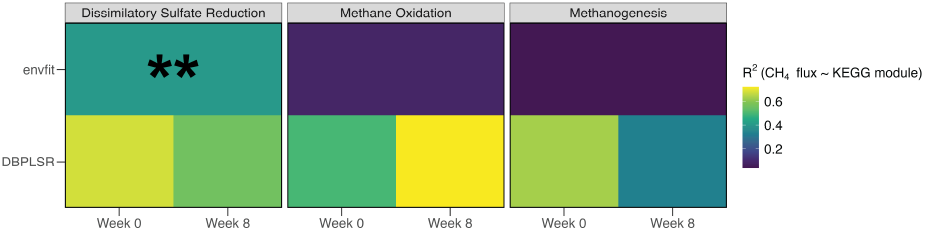
Methane flux as a function of functional gene composition. The KEGG modules analyzed were M00596 (Dissimilatory Sulfate Reduction), M00174 (Methane Oxidation), and M00617 (Methanogen). Envfit was used to assess relationships between methane (CH_4_) flux and the principal coordinate analysis (PCoA) of Kyoto Encyclopedia of Genes and Genomes (KEGG) modules, including pre- (“Week 0”) and post-experiment (“Week 8”) samples. Asterisks on envfit represent significance thresholds: ** = *p* ≤ 0.01, no label = *p >* 0.05. In the distance-based partial least squares regression (DBPLSR), week 0 and week 8 were analyzed separately. There are no p-values or significance levels for DBPLSR. Color represents the *R*^2^ for either envfit or DBPLSR (adjusted *R*^2^, two-component model for DBPLSR).

**Figure 6.**
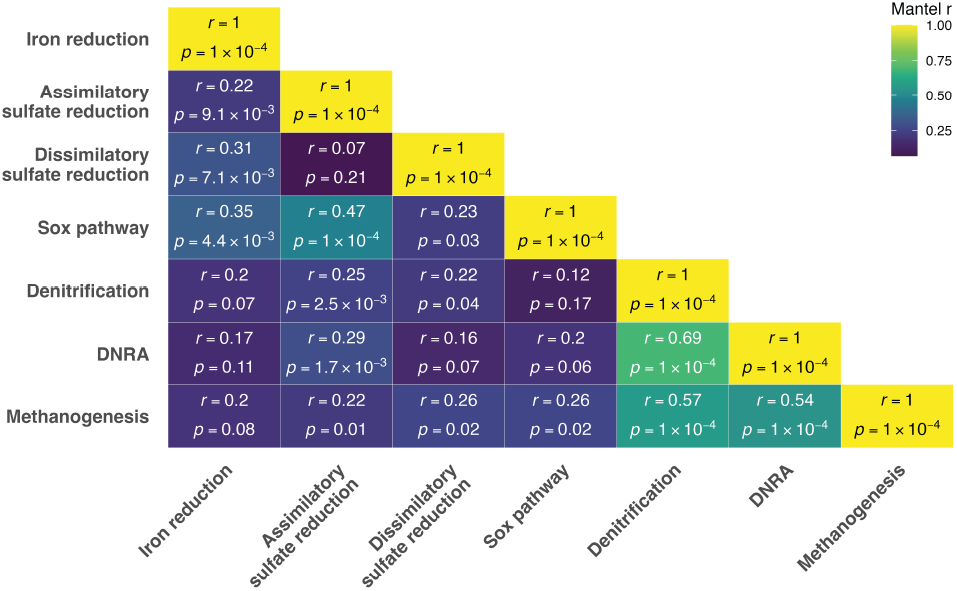
Heatmap of Mantel correlations between Bray-Curtis distance matrices of functional gene composition. Mantel tests were performed on pairs of distance matrices from samples representing wet and dry soil history, wet and dry eight-week treatment, and plant presence and absence. Correlation coefficients (*r*) and p-values are reported in the intersections between distance matrices. Warmer colors correspond to higher *r* values, while cooler colors represent lower *r* values. Abbreviations: Sox: sulfur-oxidizing multienzyme complex, DNRA: dissimilatory nitrate reduction to ammonium.

### 3.3. Iron functional genes

The composition of all Fe-active genes and the composition of Fe reduction genes were used to evaluate the response of Fe cycling communities from different hydrologic histories to flooding/drying and plant presence/absence. Soil history influenced the composition of all Fe genes identified by FeGenie (PERMANOVA: *F*_1,23_ = 4.173, *R*^2^ = 0.185, *p* = 0.031; Table 1; Table S1). The composition of Fe reduction genes identified by FeGenie also diverged according to history (PERMANOVA: *F*_1,23_ = 3.358, *R*^2^ = 0.132, *p* = 0.004; Table 1; Table S1), while the composition of Fe reduction genes based on TIGRFAMS mtrBC differed by plant presence/absence and treatment (PERMANOVA, plant: *F*_2,23_ = 9.967, *R*^2^ = 0.335, *p* = 0.002, treatment: *F*_1,23_ = 19.012, *R*^2^ = 0.319, *p* = 0.00069; Table 1; Table S1).

### 3.4. Sulfate reduction and iron reduction

We observed a significant correlation between the composition of functional genes associated with Fe reduction and assimilatory SO_4_^2−^ reduction (Mantel *r* = 0.22, *p* = 9.1 *×* 10^−3^; Figure 6). Similarly, results showed a significant correlation between functional gene composition of Fe reduction and dissimilatory SO_4_^2−^ reduction (Mantel *r* = 0.31, *p* = 7.1 *×* 10^−3^; Figure 6). Then, we assessed correlations between SO_4_^2−^ and Fe reduction under different hydrologic conditions.

Dissimilatory SO_4_^2−^ reduction and Fe reduction gene compositions were correlated in wet history (*r* = 0.65, *p* = 0.02) and wet treatment (*r* = 0.48, *p* = 0.03; Figure S5). Assimilatory SO_4_^2−^ reduction and Fe reduction gene compositions were correlated in wet treatment (*r* = 0.75, *p* = 2.3 *×* 10^−3^; Figure S5).

### 3.5. Reductive pathways linked with thiosulfate oxidation

To further evaluate redox pathways that link S, Fe, and N cycling, we measured the extent to which the metabolic gene composition of thiosulfate oxidation related to the composition of genes involved in Fe reduction, SO_4_^2−^ reduction, and NO_3_ ^−^ reduction. Based on a cutoff of *α* = 0.05, the compositions of DNRA and denitrification genes were not significantly correlated with thiosulfate oxidation by the SOX complex (Figure 6). But Fe reduction, assimilatory SO_4_^2−^ reduction, and dissimilatory SO_4_^2−^ reduction were correlated with thiosulfate oxidation to varying degrees (Figure 6).

### 3.6. Metagenome bins

Binning methods in the IMG pipeline resulted in 14 medium-to high-quality metagenome bins (Table S3). Except for one bin attributed to an Archaeal lineage (*Nitrosotalea*), all other bins were identified as bacterial taxa (Table S3). The bins were sourced from varying hydrologic histories, hydrologic treatments, and plant presence/absence treatments. We found genes for dissimilatory SO_4_^2−^ reduction and/or Fe reduction in all but four of the metagenome bins (Figure S6). Bins corresponding to Burkholderiaceae bacterium JOSHI-001 were found in both wet anddry history (Figure S6). The JOSHI-001 bin from dry history had a more complete dissimilatory SO_4_^2−^ reduction pathway than the JOSHI-001 bin from wet history (Figure S6). But the historically wet JOSHI-001 bin had both *mtrB* and *mtrC* (Fe reduction), while the historically dry JOSHI-001 bin only contained one copy of *mtrC* (Figure S6).

We also surveyed the metagenome bins for functional genes related to methanogenesis and CH_4_ oxidation. Hits for at least one methanogenic functional gene were found in all 14 metagenome bins, but only one hit for a CH_4_ oxidation gene (*pmoA-amoA*) was found in the *Nitrosotalea* bin (Figure S6).

## 4. Discussion

The impacts of SWISLR on coastal freshwater wetlands are mediated by wetland soil Fe availability, hydrologic conditions, and plant-microbe interactions (Ardón et al., 2013; Schoepfer et al., 2014; Herbert et al., 2015). In this study, we used a mesocosm approach to examine the effects of hydrologic manipulation (i.e., treatment) and plant presence/absence on soils sourced from varying histories (wet and dry). We found that soil history (wet vs dry) influenced the composition of genes involved in dissimilatory SO_4_^2−^ reduction, thiosulfate oxidation, and Fe reduction. The eight-week hydrologic treatment also modified the community composition of assimilatory SO_4_^2−^ reduction genes and resulted in correlated SO_4_^2−^ reduction and Fe reduction gene compositions, especially in the wet soil history and wet hydrologic treatment. Dissimilatory SO_4_^2−^ reduction gene composition was correlated with methanogenesis gene composition and explained more variance in CH_4_ fluxes than did methanogenesis genes. These results indicate that historical conditions strongly influenced the magnitude of soil microbial community responses to contemporary changes in hydrologic conditions and plant cover.

### 4.1. Past and current hydrologic conditions influence sulfur functional genes

The influence of hydrologic conditions, both historical and during the eight-week contemporary hydrologic treatments, was evident in the composition of S-related gene modules. The community-level compositions of SO_4_^2−^-S assimilation and assimilatory SO_4_^2−^ reduction genes both differed by hydrologic treatment, while the compositions of dissimilatory SO_4_^2−^ reduction and thiosulfate oxidation by the SOX complex both differed by soil history (Table 1; Table S1). The composition of SO_4_^2−^ reduction functional genes (assimilatory and dissimilatory) correlated with percent soil moisture (Figure 4C). This relationship revealed the importance of hydrology in the composition of SO_4_^2−^ reduction genes.

The influence of soil history on S metabolism in this coastal wetland soil is potentially two-fold: the introduction of SO_4_^2−^ via saltwater intrusion, and the reducing conditions found in waterlogged soils (in wet history), compared to more oxidizing conditions found in drier conditions (at a greater elevation above the water table). In previous work, Ardón et al. (2013) found evidence of saltwater intrusion via surface water at TOWeR. There is potential for drought-induced saltwater intrusion at this site, where the surrounding estuary’s salinity increases during drought, and winds or tides transport the brackish water upstream to the TOWeR site (Ardón et al., 2013). Flooding during storm surges is another mechanism by which surface water can facilitate saltwater intrusion (Klassen and Allen, 2017). Saltwater intrusion via either mechanism, drought or storm surge, deposits SO_4_^2−^ in the soil (Ardón et al., 2013; Schoepfer et al., 2014). This SO_4_^2−^ influx could persist in the soil and be internally transformed to other S species (sulfite, thiosulfate, sulfide, S^0^) (Ghosh and Dam, 2009; Schoepfer et al., 2014; Rückert, 2016).

Because the S from saltwater can persist in the soil, changes in soil O_2_ availability due to precipitation or dry down are important for structuring anaerobic and aerobic S metabolisms following the saltwater intrusion event (Schoepfer et al., 2014). Assimilatory SO_4_^2−^ reduction is a highly conserved process, used by both aerobic and anaerobic organisms (Rückert, 2016). On the other hand, a subset of anaerobic microorganisms participates in dissimilatory SO_4_^2−^ reduction (Rückert, 2016). Both assimilatory and dissimilatory SO_4_^2−^ reduction produce hydrogen sulfide (H_2_S) (Rückert, 2016). In assimilatory SO_4_^2−^ reduction, H_2_S is produced as an intermediate that is incorporated into biomolecules (Rückert, 2016). In contrast, dissimilatory SO_4_^2−^ reduction produces H_2_S as a waste product, in larger quantities than assimilatory SO_4_^2−^ reduction, and is therefore, a more important metabolism to consider during freshwater ecosystem sulfidization events (Hopfensperger et al., 2014; Schoepfer et al., 2014; Rückert, 2016).

While SO_4_^2−^ reduction described above consumes SO_4_^2−^, thiosulfate oxidation by the SOX complex produces SO_4_^2−^ (Ghosh and Dam, 2009; Rückert, 2016). Phototrophs, mixotrophs, and heterotrophs participate in thiosulfate oxidation via the SOX complex in a variety of assimilatory, dissimilatory, aerobic, and anaerobic pathways (Ghosh and Dam, 2009). In the context of a microbial community, thiosulfate oxidation (and S oxidation pathways in general) is important for regenerating oxidized S species (i.e., SO_4_^2−^) that can again be reduced by SRM (Ghosh and Dam, 2009). In the current study, soil history influenced the composition of thiosulfate oxidation by SOX complex genes; however, it is unclear which historical conditions (i.e., wet or dry) favored thiosulfate oxidation, as the pathway can operate in both oxic and anoxic conditions (Ding et al., 2023).

A variety of electron acceptors can be used in the oxidation of thiosulfate (Ghosh and Dam, 2009). In freshwater wetlands in Michigan, NO_3_ ^−^ reduction contributed to SO_4_^2−^ production via oxidation of sulfide (Burgin and Hamilton, 2008). Of the two major NO_3_ ^−^ reduction pathways, denitrification and DNRA, S-driven denitrification made up a greater fraction of NO_3_ ^−^ removal, but S-driven DNRA also contributed significantly to NO_3_ ^−^ removal (Burgin and Hamilton, 2008). In the estuarine landscape, sulfide oxidation can increase sediment O_2_ demand, and sediment O_2_ demand was found to be positively correlated with N_2_ flux (primarily denitrification with low potential DNRA) (Smyth et al., 2013). To test for NO_3_ ^−^-driven SO_4_^2−^ production in our mesocosm experiment, we measured correlations between the Bray-Curtis distance matrices of denitrification genes and thiosulfate oxidation genes, and DNRA genes and thiosulfate oxidation genes. Thiosulfate oxidation was not significantly correlated with either NO_3_ ^−^ reduction gene module but was correlated with Fe and SO_4_^2−^ reduction (Figure 6). The relationship between thiosulfate oxidation and SO_4_^2−^ reduction may be explained by co-occurrences of the gene modules in individual genomes, or by a diverse S cycling community that covaries with soil S concentrations. *Sox* and *dsr* genes co-occur in multiple species and may be expressed simultaneously, although *dsr* is likely used in sulfide oxidation in these cases (Beller et al., 2006; Grimm et al., 2011). These genome co-occurrences could contribute to the correlations we observed between thiosulfate oxidation and SO_4_^2−^ reduction genes, but the community-wide relationship is likely driven by multiple species. A quantitative survey of *sox* and *dsr* genes in an estuarine salt marsh found high abundances of both genes across a tidal gradient, and inferred that sulfide oxidation and dissimilatory SO_4_^2−^ reduction work in concert to recycle sulfide and SO_4_^2−^ to sustain sulfide levels that are tolerable to the plant community, and SO_4_^2−^ levels to sustain heterotrophic soil respiration of plant C inputs (Zheng et al., 2017). It was also found that Fe(III) was the primary driver of the community compositions of SRM and FeRM (Zheng et al., 2017). Similarly, we found a correlation between genes for thiosulfate oxidation and Fe reduction (Figure 6). Again, these functional capabilities could be found within species (Kato and Ohkuma, 2021), and thiosulfate may serve as an intermediate in the direct coupling of Fe reduction with S oxidation (Breuker and Schippers, 2024). This relationship, in conjunction with SO_4_^2−^ reduction (discussed below), links S and Fe cycling in these Fe-rich and SWISLR-impacted coastal wetland soils.

### 4.2. Soil history influences iron functional gene composition

Soil history modified the Fe functional gene composition. Specifically, soil history altered the composition of all Fe genes identified by FeGenie and the subset of Fe reduction genes identified by FeGenie (Table 1; Table S1). Transformations of Fe vary based on the redox status and pH of the environment (Weber et al., 2006). It follows that dry or wet, and oxic or anoxic historical conditions would structure different Fe cycling communities. In anoxic environments (with pH *>* 4), microbial Fe(III) reduction is a major electron sink for organic matter oxidation (Canfield et al., 1993; Weber et al., 2006). Therefore, greater genetic potential for Fe reduction may exist in historically wet soils with presumably lower O_2_ availability. But this inference is complicated by the ongoing microbial transformations of N alongside aerobic and anaerobic oxidations and reductions of Fe (reviewed in Weber et al., 2006). Our analysis of the composition of all Fe genes identified by FeGenie highlights the impact of soil history on Fe cycling and Fe reduction specifically.

The FeGenie tool was developed in response to a lack of Fe annotation tools (Garber et al., 2020). Before FeGenie, one of the few options for identifying Fe reduction genes was TIGRFAMS *mtrB* (TIGR03509) and *mtrC* (TIGR03507) (Garber et al., 2020). We also used the composition of TIGR03509 and TIGR03507 to analyze genetic potential for Fe reduction and found the composition to differ by plant (presence/absence) and hydrologic treatment (Table 1; Table S1). The genes *mtrB* and *mtrC* are included in FeGenie, but FeGenie and IMG’s TIGRFAM annotation pipeline resulted in different counts of *mtrB* and *mtrC*. These contrasting results from different annotation tools highlight the current uncertainties in Fe gene annotation and point towards a need for increased consensus in Fe gene annotation.

### 4.3. Sulfate reduction and iron reduction correlate in reducing conditions

We hypothesized that SO_4_^2−^ reduction and Fe reduction would be coupled (meaning significantly correlated) in reducing (i.e., flooded) conditions. We found that the community compositions of SO_4_^2−^ reduction and Fe reduction genes were significantly correlated across soil histories, hydrologic treatments, and plant presence/absence (Figure 6). When we measured the relationship within soil histories and hydrologic treatments, we found SO_4_^2−^ and Fe reduction correlated in wet conditions (history and treatment) and not in dry conditions (Figure S5). This indicates that the functional gene compositions of both metabolisms were shaped by long-term hydrologic regimes in the field and responded similarly and strongly to eight weeks of flooding. Because the association was based on DNA sequencing, it indicates that the relative abundance of SRM and FeRM changed in response to hydrologic treatment over eight weeks of incubation but does not reflect the rates of SO_4_^2−^ reduction and Fe reduction. Both SO_4_^2−^ reduction and Fe reduction are commonly found in anoxic sediments (Flynn et al., 2021). The wet treatment appears to have imposed suboxic/anoxic conditions, characterized by a higher percent of paint removed from IRIS tubes (Figure S2), and these reducing conditions were likely conducive to both SRM and FeRM. This supports previous findings of correlated SO_4_^2−^ and Fe reduction in TOWeR soils (Schoepfer et al., 2014).

### 4.4. Dissimilatory sulfate reduction gene composition explains variance in methane fluxes

In addition to the Fe-S linkages described above, dissimilatory SO_4_^2−^ reduction gene composition was related to the production of CH_4_. Dissimilatory SO_4_^2−^ reduction gene composition was significantly correlated with CH_4_ fluxes (Figures 3C and 5). At the start of the experiment (week zero), the composition of dissimilatory SO_4_^2−^ reduction genes had a stronger relationship with CH_4_ fluxes than either CH_4_ oxidation or methanogenesis functional genes (Figure 5, Table S2). At the end of the experiment, dissimilatory SO_4_^2−^ reduction still explained a greater percentage of variance in CH_4_ fluxes than methanogenesis, but less than CH_4_ oxidation (Figure 5, Table S2). The CH_4_ oxidation gene composition increased in importance from week zero to week eight, but no treatment impacts on CH_4_ oxidation gene composition were observed (Table 1). During the first GHG time point (June 13th), CH_4_ fluxes were greatest in wet soil history, followed by interim history, but the impact of history on CH_4_ fluxes abated thereafter (Figure S3). This contrasts with a drying-rewetting microcosm study that found greater potential for methanogenesis upon rewetting in soils from ephemeral wetlands vs. standing water sites (Kannenberg et al., 2015). This is perhaps explained by our initial GHG sampling occurring two weeks after the initiation of hydrologic treatments, and our inclusion of plants to replenish soil C substrates. The methanogen populations were shaped by historical conditions but responded to hydrologic treatments throughout the middle three time points (Figure S4). Methanogenic functional gene abundance did not reflect differences in hydrologic treatment Bledsoe et al. (2025), indicating changes to gene expression and protein activity were used to respond to hydrologic treatments, with little detectable change to methanogenic gene abundance. The strength of the relationship between CH_4_ fluxes and dissimilatory SO_4_^2−^ reduction genes mirrored that of methanogen gene composition, dropping from week zero to week eight, but was higher for dissimilatory SO_4_^2−^ reduction at both time points (Figure 5). This caused us to question whether the relationship between methanogens and SRM was mutualistic, competitive, or covarying with hydrologic conditions.

Our compositional approach showed that variation in dissimilatory SO_4_^2−^ reduction gene composition corresponded with variance in CH_4_ fluxes and methanogen functional gene composition. A weak positive correlation between the distance matrices of methanogenic and dissimilatory sulfate-reducing functional genes indicated that samples with similar communities of methanogens also tended to have similar communities of SO_4_^2−^ reducers. While Fe reduction gene composition was not strongly related to CH_4_ fluxes, it was correlated with SO_4_^2−^ reduction gene composition (Figure 6). Fe reduction is known to inhibit methanogenesis through substrate competition (acetate and hydrogen), both between separate groups of FeRM and methanogens capable of Fe reduction (van Bodegom et al., 2004; Sivan et al., 2016). Ferric iron compounds have also been shown to directly inhibit methanogenesis (van Bodegom et al., 2004). The interactions among dissimilatory SO_4_^2−^ reduction, Fe reduction, and methanogenesis are complex and may structure ecosystem-level transformations of S, Fe, and C.

Explanations for both positive and negative relationships between SO_4_^2−^ and Fe reduction genes and CH_4_ fluxes are possible. Dissimilatory SO_4_^2−^ reduction, Fe reduction, and methanogenesis are all generally considered anaerobic metabolisms (Lyu et al., 2018; Flynn et al., 2021). Therefore, the reducing or oxidizing environment (e.g., wet vs. dry conditions) could have impacted all three metabolisms similarly, resulting in the observed correlations. Alternatively, the correlation between dissimilatory SO_4_^2−^ reduction and CH_4_ fluxes could be the result of competition between microbial taxa. Since Fe reduction and dissimilatory SO_4_^2−^ reduction are more thermodynamically favorable than methanogenesis (in standard conditions) (Wang et al., 2017), it is conceivable that when FeRM and SRM are abundant (because Fe(III) and SO_4_^2−^ are available), these microbial communities outcompete the less abundant methanogens, resulting in a negative correlation. Similarly, methanogenic archaea may switch from methanogenesis to Fe reduction when resources allow (Sivan et al., 2016), and the presence of Fe reduction and methanogenic functional genes in three metagenome bins indicates genetic potential for this switch, but none of the bins were assigned to methanogenic lineages (Figure S6). The specific resources that methanogens and SRM compete for are acetate and hydrogen (Schink, 1997; Conrad, 1999; Chidthaisong and Conrad, 2000). Methylotrophic methanogens avoid competition with SRM by consuming methyl compounds that are not consumed by SRM (Oremland and Polcin, 1982; Kiene et al., 1986). Bueno de Mesquita et al. (2024) incubated TOWeR soils with artificial seawater, with and without SO_4_^2−^, but found no increase in methyl-reducing methanogens, indicating that avoiding competition with SRM during a pulse of SO_4_^2−^ availability was not a widely adopted strategy by the TOWeR methanogen community. In addition to substrate competition, SO_4_^2−^ reduction may directly inhibit methanogenesis through the production of sulfide, but some methanogens can tolerate high sulfide concentrations (Bryant et al., 1977; Mountfort and Asher, 1979; Koster et al., 1986). Given that dissimilatory SO_4_^2−^ reduction and methanogenesis functional genes were positively correlated (Figure 6, Figure S5) and both decreased in their ability to explain variance in CH_4_ fluxes from start to end of the experiment (Figure 5), competition between the two functional groups is not well supported by our data. Instead, mutualism may explain the concomitant shifts in variance explained in CH_4_ fluxes and the strong relationship between dissimilatory SO_4_^2−^ reduction and CH_4_ fluxes (Figures 3C, 6). A known mutualism between SRM and methanogens exists, in which methanogens consume H_2_S produced by SRM, and thereby maintain the required pH for both SO_4_^2−^ reduction and methanogenesis to proceed (Shi et al., 2020). Relationships between SRM and methanogens vary between environments and experimental setups and are commonly tied to C substrate concentrations (Hershey et al., 2014; Kannenberg et al., 2015; Bueno de Mesquita et al., 2024). Our functional gene analysis based on metagenomic sequencing and *in situ* CH_4_ fluxes supports a mutualistic, or environmentally co-varying relationship between SRM and methanogens. To consistently predict freshwater wetland CH_4_ emissions under future SWISLR, experiments that isolate the impact of SO_4_^2−^ on CH_4_ emissions (like Bueno de Mesquita et al. (2024)), will need to be replicated in a range of freshwater wetland soils with the inclusion of wetland plants to regulate C and O_2_ availability.

## 5. Conclusions

An oversimplified view of biogeochemistry might describe the biogeochemical cycles of C, N, Fe, and S in isolation, with transformations structured in communities according to thermodynamic favorability (Bethke et al., 2011; Schlesinger et al., 2011). But it is increasingly clear that biogeochemical cycles are intertwined, and a “microbial energy economy” that relies on resource availability and biotic and abiotic interactions is closer to reality than a hierarchically structured thermodynamic ladder (Bethke et al., 2011; Burgin et al., 2011). The specific biotic and abiotic context in which communities are found will likely determine which linkages between cycles are relevant. In the wetland soil mesocosm study presented here, Fe-rich soils and seasonal saltwater intrusion importing SO_4_^2−^ make Fe and S linkages with C and N metabolism particularly relevant. Considering linked biogeochemical cycles and the historical context-dependence of an ecosystem will help improve predictions of the future impacts of SWISLR on coastal biogeochemistry.

## 6 Contributions

Contributed to conception and design: CGF, ALP Contributed to acquisition of data: ALP

Contributed to analysis and interpretation of data: CGF, ALP

Drafted and/or revised the article: CGF, ALP

Approved the submitted version for publication: CGF, ALP

## 7. Acknowledgments

Thank you to R. Bledsoe for developing and executing the experiment. We extend our gratitude to J. LeCrone, L. Armstrong, M. Stillwagon, G. Gunderson, C. Eakins, J. Basco, and C. Bledsoe for their invaluable assistance in the field and laboratory. We also appreciate J. Gill and the East Carolina University grounds crew for their efforts in maintaining the area around the shade house. Additionally, we thank M. Muscarella for contributing to the microbiome analyses. Thank you to J. Hoben and E. Field for manuscript revisions and analysis ideas. E. Field and C. Joyner helped with FeGenie troubleshooting and shared computing resources.

## 10. Supplemental material

The supplemental files for this article can be found in the file: Supplemental Material.pdf

## 11. Data accessibility statement

All code and data used in this study are in a public GitHub repository (https://github.com/PeraltaLab/WetlandMesocosm_GHG_Timberlake) with a Zenodo DOI (10.5281/zenodo.4042109). Raw amplicon sequence files can be found at NCBI SRA BioProject ID PRJNA636184, and metagenome sequence files can be found at NCBI SRA BioProject ID PRJNA641216.

