## Supplemental_Material for "Shifting flood regimes alter iron–sulfur metabolism and greenhouse gas associations"

for

Publication in Elementa—Science of the Anthropocene

8 October 2025

**Contents**

Figure S1. Conceptual diagram depicting relative ranges of redox potentials associated with predicted microbial processes ………………………… page number 3

Figure S2. Percentage of paint removed from IRIS tubes during mesocosm incubation ………………………...page number 4

Figure S3. Differences in greenhouse gas fluxes across hydrologic histories ……………………………………………………page number 5

Figure S4. Greenhouse gas fluxes measured under different hydrologic treatments over time …………………………………………………… page number 6

Figure S5. Hydrology-specific Mantel heatmap of correlations between Bray-Curtis distance matrices of functional gene modules ..………………….. page number 8

Figure S6. Heatmap of functional genes found in metagenome bins ……page number 9

Table S1. Summary of permutational multivariate analysis of variance (PERMANOVA) comparing microbial community composition due to main effects ………page number 10

Table S2. Summary of distance-based partial least squares regression representing the percent of variance in greenhouse gas fluxes explained by the first two components of models derived from functional gene composition …………page number 12

Table S3. Summary of metagenome bins …………………………………page number 14

**FIGURES**
**Figure S1**

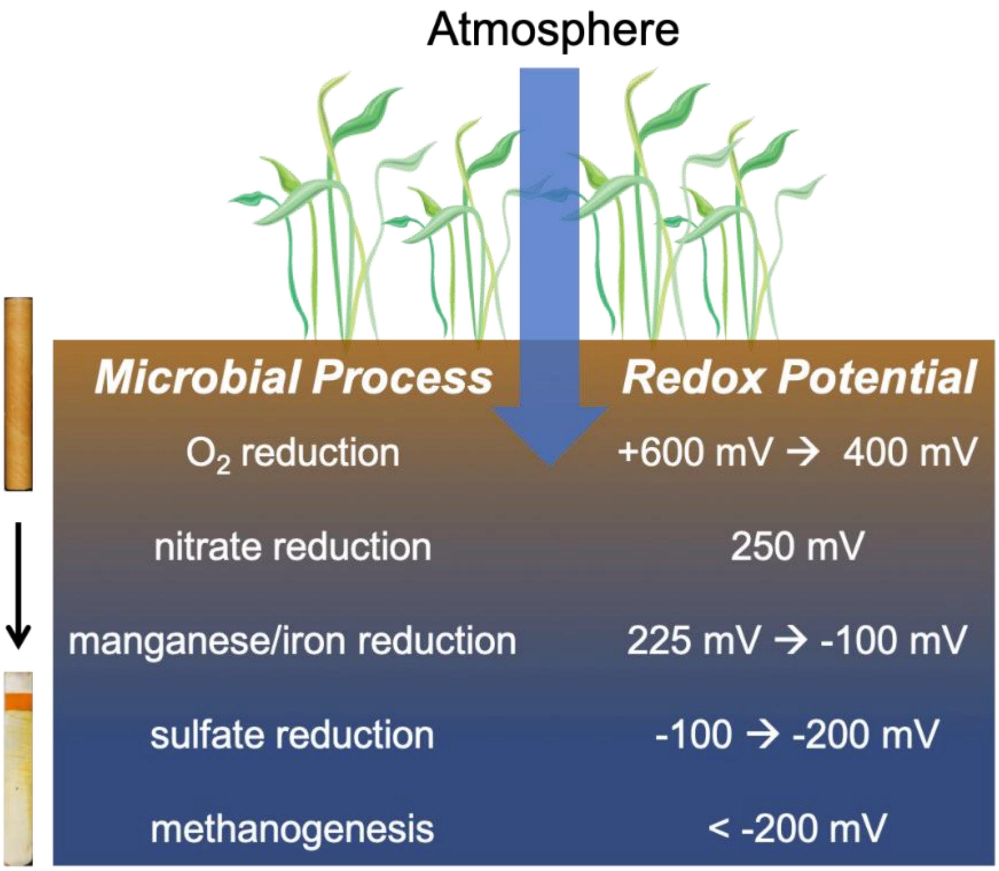


**Figure S1. Conceptual diagram depicting relative ranges of redox potentials associated with predicted microbial processes.** On the left side of the diagram, indicator for reduction in soils (IRIS) tubes are used to measure soil redox status, where iron oxide paint in orange represents oxidized (Fe(III)) conditions, and white represents reduced (Fe(II)) conditions.

**Figure S2**

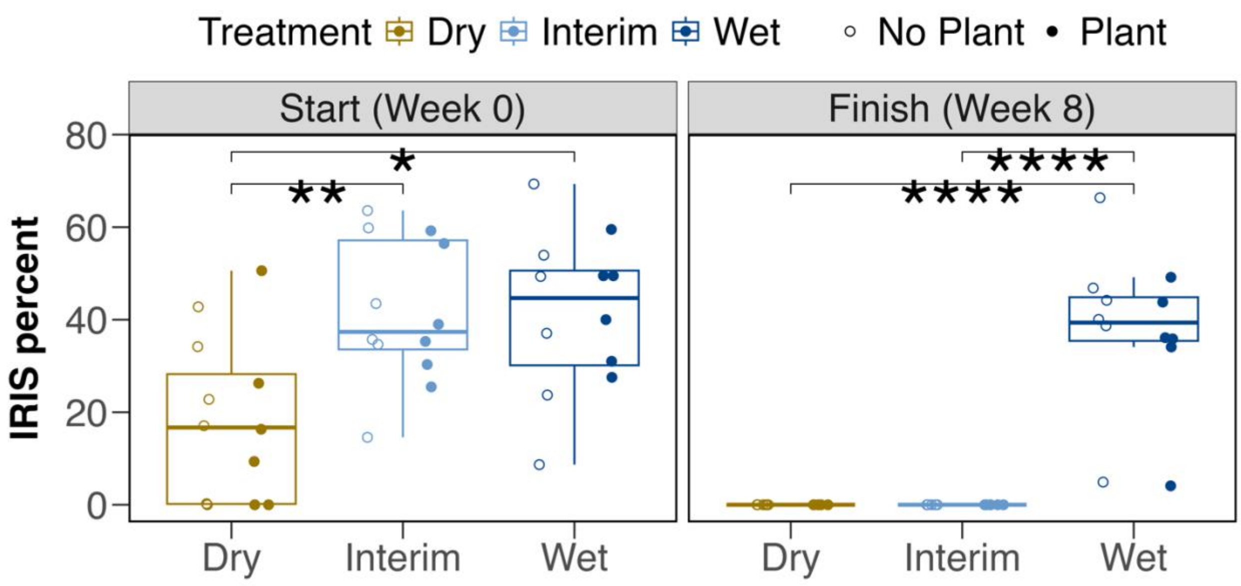


**Figure S2. Percentage of paint removed from IRIS tubes during mesocosm incubation.** More reducing conditions correspond to a higher percent of paint removed from IRIS tubes ("IRIS percent"). Individual data points from IRIS tubes are plotted as points; the symbol color represents the hydrologic treatment, with open points indicating no plant and closed points indicating a plant. The boxplot is a visual representation of the following summary statistics: the median, the 25$^{th}$ and 75$^{th}$ percentiles, and the whiskers, which represent 1.5 × the interquartile range. Asterisks and brackets represent significance levels of pairwise comparisons using the Dwass-Steel-Critchlow-Fligner procedure: ****: $p \leq 0.0001$, **: $p \leq 0.01$, *: $p \leq 0.05$, no asterisk: nonsignificant. Figure modified from Bledsoe et al. (2025).

**Figure S3**

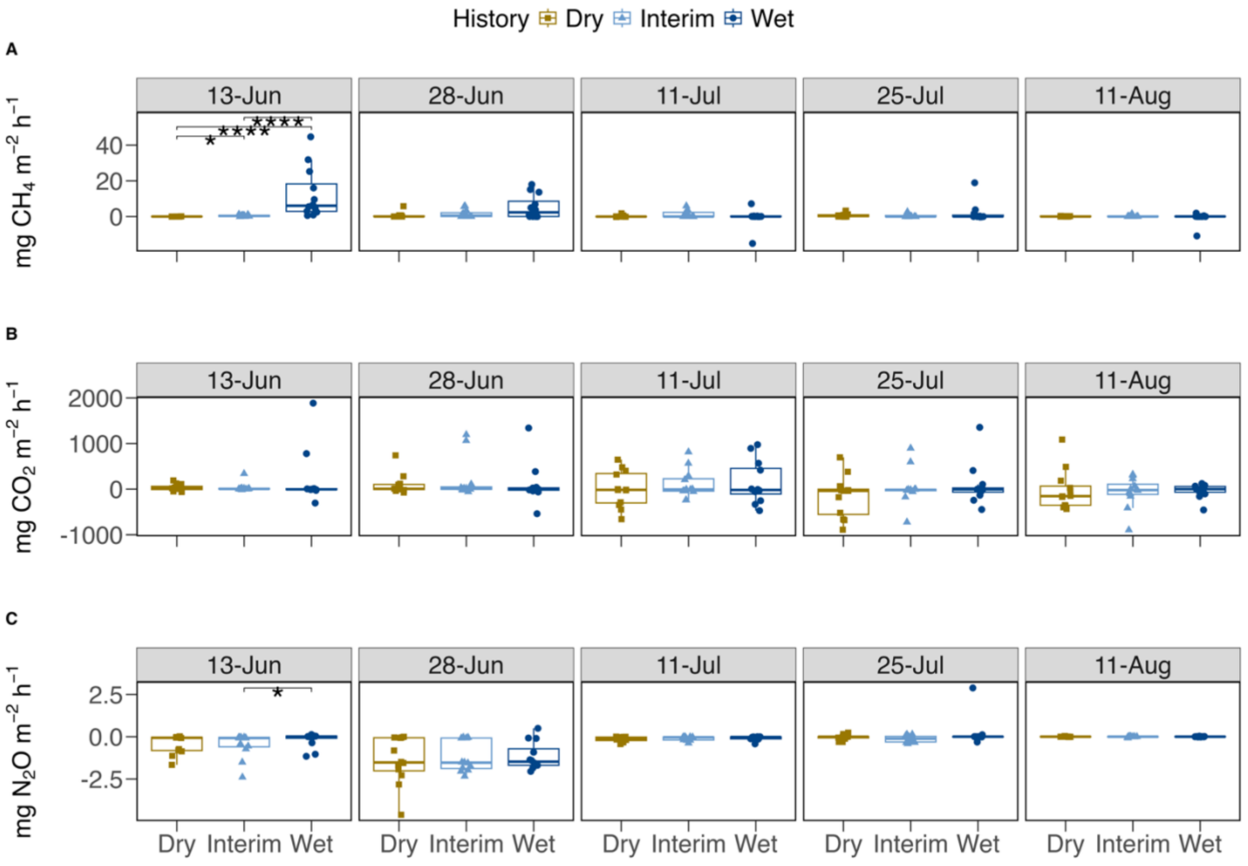


**Figure S3. Differences in greenhouse gas fluxes across hydrologic histories.** Fluxes are in milligrams of greenhouse gas per square meter per hour. Individual fluxes are plotted as points. Colors and shapes represent soil history (brown square = dry, light blue triangle = interim, dark blue circle = wet). Boxplots summarize median, first and third quartiles, and two whiskers extending |≤| 1.5 * interquartile range. Asterisks and brackets represent significance levels of pairwise comparisons using the Dwass-Steel-Critchlow-Fligner procedure: ****: $p \leq 0.0001$, *: $p \leq 0.05$, no asterisk: nonsignificant. An extreme outlier of 248 mg $CH_4$ $m^{-2}$ $h^{-1}$ on July 11th, measured in a mesocosm sourced from interim history, was used in statistical calculations but not displayed in the plot to improve visualization of differences across treatments.

#### Figure S4

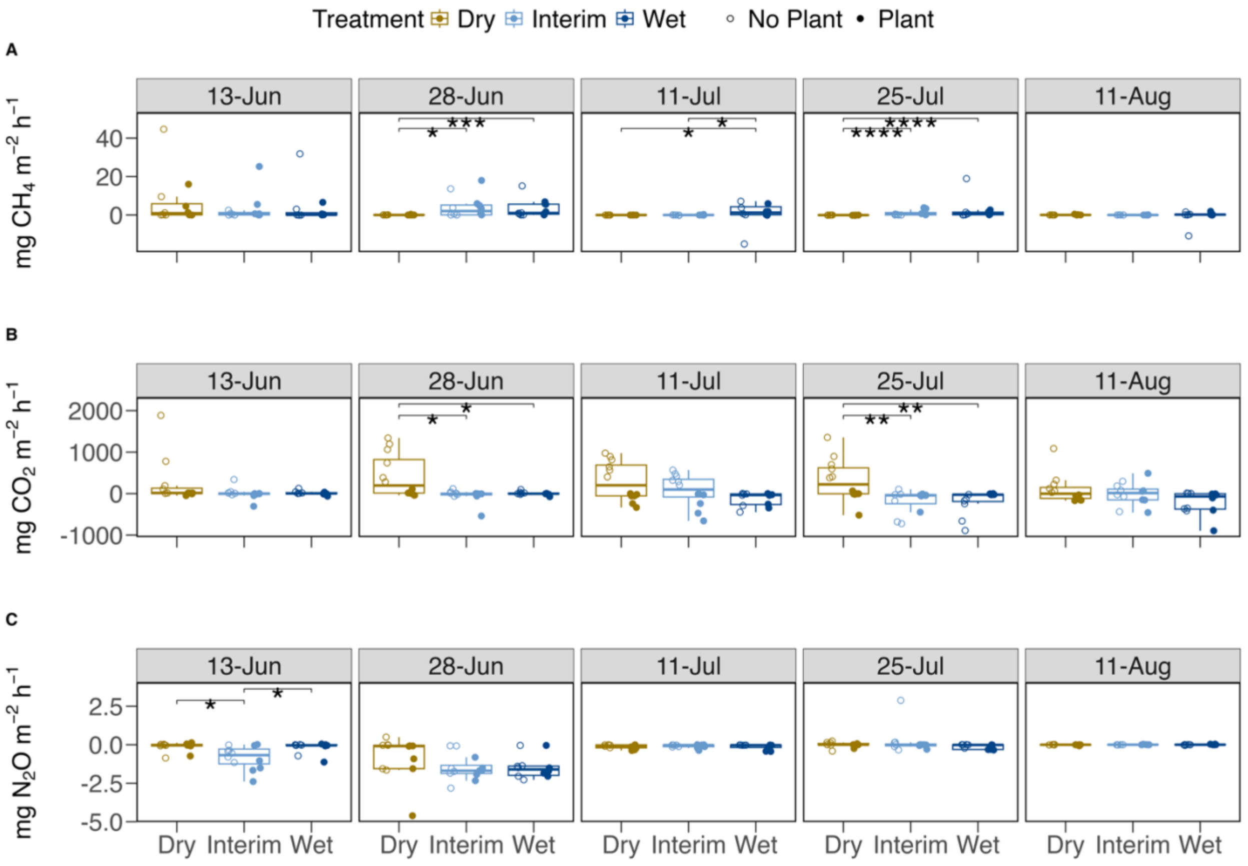


**Figure S4. Greenhouse gas fluxes measured under different hydrologic treatments over time.** Fluxes are in milligrams of greenhouse gas per square meter per hour. Color represents hydrologic treatment (brown = dry, light blue = interim, dark blue = wet). Individual fluxes are plotted, and fill represents plant presence/absence: open point = no plant, filled point = plant. Boxplots summarize median, first and third quartiles, and two whiskers extending |≤| 1.5 * interquartile range. Asterisks and brackets represent significance levels of pairwise comparisons among hydrologic treatments using the Dwass-Steel-Critchlow-Fligner procedure: ****: $p \leq 0.0001$, ***: $p \leq 0.001$, **: $p \leq 0.01$, *: $p \leq 0.05$, no asterisk: nonsignificant. An extreme outlier of 248 mg $CH_4$ $m^{-2}$ $h^{-1}$, measured in a wet treatment with no plants on July 11th, was used in statistical calculations but not displayed in the plot to improve visualization of differences across treatments.

#### Figure S5

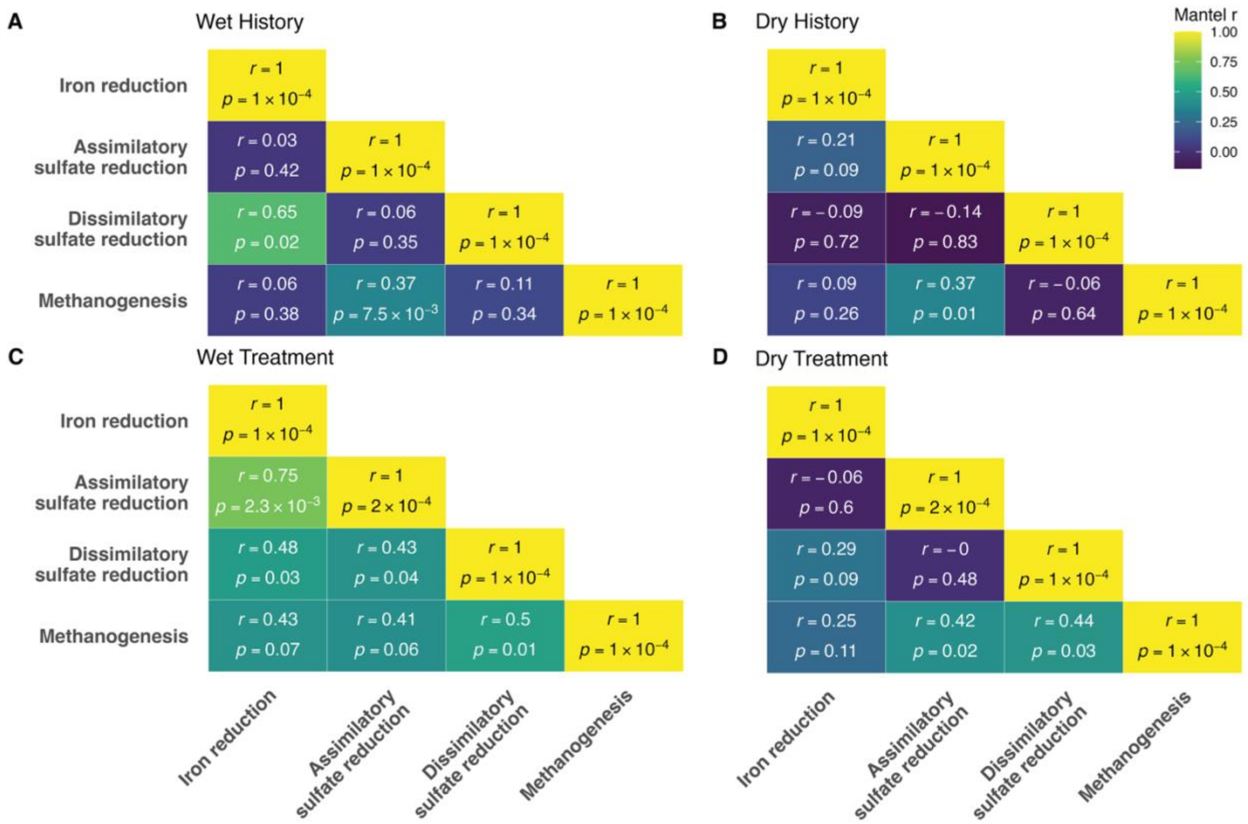


**Figure S5. Hydrology-specific Mantel heatmap of correlations between Bray-Curtis distance matrices of functional gene modules.** Distance matrices were subset by soil history (wet vs. dry) and hydrologic treatment (wet vs. dry), and Mantel tests were performed on pairs of distance matrices within each subset. Correlation coefficients (*r*) and *p*-values are reported in the intersections between distance matrices. Warmer colors correspond to higher *r* values, while cooler colors represent lower *r* values.

**Figure S6**

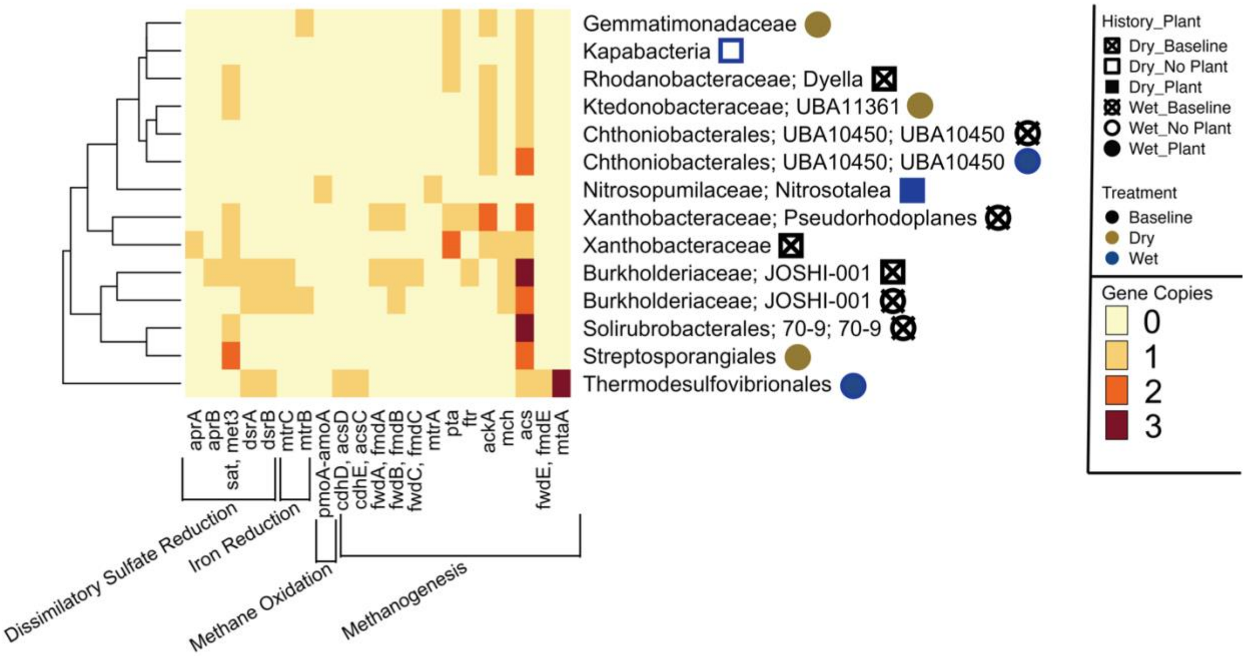


**Figure S6. Heatmap of functional genes found in metagenome bins.** Symbols next to bin taxonomy show the environment the bin was sourced from: colors refer to hydrologic treatments (brown = dry, dark blue = wet), shapes refer to soil history (square = dry, circle = wet), and the shape fill represents plant treatment (open symbol = no plant, closed symbol = plant). Heatmap colors indicate the number of gene copies found in a metagenome bin. The dendrogram on the left side is calculated based on the similarity of the gene counts in the functional modules listed at the bottom of the heatmap. The functional gene modules are based on KEGG Dissimilatory Sulfate Reduction (M00596), TIGRFAMS mtrBC (TIGR03509, TIGR03507), KEGG Methane Oxidation (M00174), and KEGG Methanogen (M00617). The complete set of genes in each module was queried, but only genes with at least one hit in one bin were included in the heatmap.

### TABLES

#### Table S1. Summary of permutational multivariate analysis of variance (PERMANOVA) comparing microbial community composition due to main effects.

| Gene Module | Main Effect | DF[a] | SumSq[b] | $R^2$ | *F* | *p* |
|---|---|---|---|---|---|---|
| Sulfate-Sulfur Assimilation (KEGG[c] M00616) | Plant | 2 | 0.006 | 0.066 | 0.891 | 0.527 |
| | History | 1 | 0.005 | 0.061 | 1.654 | 0.153 |
| | **Treatment[d]** | 1 | **0.013** | **0.144** | **3.917** | **0.008** |
| | Plant × History | 2 | 0.008 | 0.090 | 1.228 | 0.276 |
| | Plant × Treatment | 1 | 0.004 | 0.051 | 1.389 | 0.222 |
| Assimilatory Sulfate Reduction (KEGG M00176) | Plant | 2 | 0.006 | 0.066 | 0.872 | 0.514 |
| | History | 1 | 0.004 | 0.046 | 1.221 | 0.288 |
| | **Treatment** | 1 | **0.015** | **0.156** | **4.111** | **0.013** |
| | Plant × History | 2 | 0.009 | 0.091 | 1.193 | 0.307 |
| | Plant × Treatment | 1 | 0.003 | 0.031 | 0.817 | 0.497 |
| Dissimilatory Sulfate Reduction (KEGG M00596) | Plant | 2 | 0.012 | 0.113 | 1.859 | 0.132 |
| | **History** | 1 | **0.032** | **0.316** | **10.397** | **$3 \times 10^{-4}$** |
| | Treatment | 1 | 0.003 | 0.029 | 0.947 | 0.388 |
| | Plant × History | 2 | 0.005 | 0.046 | 0.755 | 0.577 |
| | Plant × Treatment | 1 | 0.001 | 0.010 | 0.3171 | 0.784 |
| Sulfate Reduction (combined assimilatory and dissimilatory) | Plant | 2 | 0.007 | 0.060 | 0.786 | 0.619 |
| | **History** | 1 | **0.012** | **0.108** | **2.830** | **0.033** |
| | **Treatment** | 1 | **0.013** | **0.111** | **2.906** | **0.026** |
| | Plant × History | 2 | 0.009 | 0.077 | 1.016 | 0.418 |
| | Plant × Treatment | 1 | 0.004 | 0.034 | 0.879 | 0.476 |
| Thiosulfate Oxidation by SOX Complex (KEGG M00595) | Plant | 2 | 0.004 | 0.045 | 0.608 | 0.637 |
| | **History** | 1 | **0.022** | **0.243** | **6.547** | **0.009** |
| | Treatment | 1 | 0.007 | 0.078 | 2.095 | 0.135 |
| | Plant × History | 2 | 0.002 | 0.017 | 0.234 | 0.939 |
| | Plant × Treatment | 1 | 0.002 | 0.024 | 0.656 | 0.501 |
| Fe Genes (FeGenie) | Plant | 2 | 0.001 | 0.028 | 0.315 | 0.880 |
| | **History** | 1 | **0.006** | **0.185** | **4.173** | **0.031** |
| | Treatment | 1 | 0.001 | 0.044 | 0.994 | 0.354 |
| | Plant × History | 2 | 0.001 | 0.028 | 0.318 | 0.872 |
| | Plant × Treatment | 1 | $2 \times 10^{-4}$ | 0.006 | 0.139 | 0.909 |
| Fe Reduction (FeGenie) | Plant | 2 | 0.157 | 0.094 | 1.189 | 0.280 |
| | **History** | 1 | **0.221** | **0.132** | **3.358** | **0.004** |
| | Treatment | 1 | 0.010 | 0.060 | 1.513 | 0.161 |
| | Plant × History | 2 | 0.102 | 0.061 | 0.772 | 0.700 |
| | Plant × Treatment | 1 | 0.038 | 0.023 | 0.580 | 0.787 |
| Fe Reduction (TIGRFAMS mtrBC) | **Plant** | 2 | **0.032** | **0.335** | **9.967** | **0.002** |
| | History | 1 | 0.004 | 0.037 | 2.187 | 0.162 |

| | | | | | | |
|---|---|---|---|---|---|---|
| | **Treatment** | 1 | **0.031** | **0.319** | **19.012** | **7 x 10$^{-4}$** |
| | Plant × History | 2 | 0.003 | 0.036 | 1.058 | 0.367 |
| | Plant × Treatment | 1 | 5 x 10$^{-4}$ | 0.005 | 0.288 | 0.598 |
| Dissimilatory Nitrate Reduction to Ammonium (DNRA) (KEGG M00530) | Plant | 2 | 0.014 | 0.115 | 1.282 | 0.270 |
| | History | 1 | 0.004 | 0.035 | 0.773 | 0.529 |
| | Treatment | 1 | 0.006 | 0.049 | 1.090 | 0.355 |
| | Plant × History | 2 | 0.009 | 0.079 | 0.881 | 0.518 |
| | Plant × Treatment | 1 | 0.001 | 0.007 | 0.156 | 0.951 |
| Methane Oxidation (KEGG M00174) | Plant | 2 | 0.171 | 0.085 | 0.878 | 0.542 |
| | History | 1 | 0.051 | 0.025 | 0.518 | 0.704 |
| | Treatment | 1 | 0.023 | 0.012 | 0.240 | 0.888 |
| | Plant × History | 2 | 0.129 | 0.064 | 0.663 | 0.701 |
| | Plant × Treatment | 1 | 0.083 | 0.041 | 0.851 | 0.490 |

[a]Bold text indicates significant differences ($p \leq 0.05$). These differences are summarized in Table 1.

[b]Sum of squares (SumSq)

[c]Kyoto Encyclopedia of Genes and Genomes (KEGG)

[d]Degrees of freedom (DF)

**Table S2. Summary of distance-based partial least squares regression representing the percent of variance in greenhouse gas fluxes explained by the first two components of models derived from functional gene composition.**

| $CH_4$[a] flux ~ Gene module[b] | Time point | n comp total[c] | Statistic | Comp 1 | Comp 2 |
|---|---|---|---|---|---|
| $CH_4$ flux ~ Dissimilatory Sulfate Reduction | Week 0 | 7 | $R^2$ | 0.60 | 0.77 |
| | | | adjusted $R^2$ | 0.53 | 0.68 |
| | | | gvar[d] | 67.13 | 86.25 |
| | | | crit[e] | 5.12 | 3.93 |
| $CH_4$ flux ~ Dissimilatory Sulfate Reduction | Week 8 | 15 | $R^2$ | 0.46 | 0.62 |
| | | | adjusted $R^2$ | 0.42 | 0.56 |
| | | | gvar | 53.15 | 85.41 |
| | | | crit | 0.28 | 0.23 |
| $CH_4$ flux ~ Methane Oxidation | Week 0 | 7 | $R^2$ | 0.44 | 0.64 |
| | | | adjusted $R^2$ | 0.35 | 0.50 |
| | | | gvar | 76.22 | 86.27 |
| | | | crit | 7.16 | 6.28 |
| $CH_4$ flux ~ Methane Oxidation | Week 8 | 15 | $R^2$ | 0.63 | 0.77 |
| | | | adjusted $R^2$ | 0.61 | 0.73 |
| | | | gvar | 29.28 | 84.80 |
| | | | crit | 0.19 | 0.14 |
| $CH_4$ flux ~ Methanogenesis | Week 0 | 7 | $R^2$ | 0.52 | 0.74 |
| | | | adjusted $R^2$ | 0.44 | 0.63 |
| | | | gvar | 26.04 | 62.04 |
| | | | crit | 6.10 | 4.56 |
| $CH_4$ flux ~ Methanogenesis | Week 8 | 15 | $R^2$ | 0.19 | 0.42 |
| | | | adjusted $R^2$ | 0.13 | 0.33 |
| | | | gvar | 35.78 | 58.00 |
| | | | crit | 0.43 | 0.35 |

[a]Methane ($CH_4$)

[b]Functional gene distance matrices are based on the Bray-Curtis dissimilarities of gene relative abundances within respective KEGG modules (see main text for KEGG module numbers).

[c]The total number of components in each model (n comp total) is equal to the sample size minus one.

[d]Total weighted geometric variability (gvar)

[e] generalized cross-validation critical value (crit)

**Table S3. Summary of metagenome bins.**

| Bin ID[a] | History | Treatment | Plant Presence /Absence | Bin Quality | GTDBTK Lineage | Bin Completeness | Bin Contamination | Total Bases | Gene Count |
|---|---|---|---|---|---|---|---|---|---|
| 3300036865_5 | Dry | Baseline | Baseline | MQ[b] | Bacteria; Proteobacteria; Gammaproteobacteria; Betaproteobacteriales; Burkholderiaceae; JOSHI-001 | 88.02 | 6.17 | 4674782 | 4849 |
| 3300036991_3 | Dry | Baseline | Baseline | HQ[c] | Bacteria; Proteobacteria; Gammaproteobacteria; Xanthomonadales; Rhodanobacteraceae; *Dyella* | 98.45 | 2.71 | 4930595 | 4528 |
| 3300036991_4 | Dry | Baseline | Baseline | MQ | Bacteria; Proteobacteria; Alphaproteobacteria; Rhizobiales; Xanthobacteraceae | 54.51 | 1.92 | 2325436 | 2615 |
| 3300036870_7 | Dry | Wet | Plant | MQ | Archaea; Crenarchaeota; Nitrososphaeria; Nitrososphaerales; Nitrosopumilaceae; *Nitrosotalea* | 61.00 | 0.97 | 780222 | 1032 |
| 3300036873_3 | Dry | Wet | No Plant | HQ | Bacteria; Bacteroidota; Kapabacteria | 96.98 | 1.64 | 3157872 | 2807 |
| 3300036867_3 | Wet | Baseline | Baseline | MQ | Bacteria; Proteobacteria; Gammaproteobacteria; Betaproteobacteriales; Burkholderiaceae; JOSHI-001 | 76.88 | 0.70 | 3880859 | 4065 |
| 3300036989_2 | Wet | Baseline | Baseline | MQ | Bacteria; Proteobacteria; Alphaproteobacteria; Rhizobiales; | 62.92 | 8.67 | 3626868 | 4149 |

| | | | | | | | | | |
|---|---|---|---|---|---|---|---|---|---|
| | | | | | Xanthobacteraceae; *Pseudorhodoplanes* | | | | |
| 3300036989_4 | Wet | Baseline | Baseline | MQ | Bacteria; Actinobacteriota; Thermoleophilia; Solirubrobacterales; 70-9; 70-9 | 80.23 | 5.11 | 2422891 | 2808 |
| 3300036990_3 | Wet | Baseline | Baseline | MQ | Bacteria; Verrucomicrobiota; Verrucomicrobiae; Chthoniobacterales; UBA10450; UBA10450 | 61.40 | 0.72 | 2180017 | 2388 |
| 3300036875_5 | Wet | Dry | Plant | MQ | Bacteria; Actinobacteriota; Actinobacteria; Streptosporangiales | 51.10 | 8.78 | 3331403 | 3616 |
| 3300036898_4 | Wet | Dry | Plant | MQ | Bacteria; Chloroflexota; Ktedonobacteria; Ktedonobacterales; Ktedonobacteraceae; UBA11361 | 85.26 | 1.98 | 4109904 | 4252 |
| 3300036898_5 | Wet | Dry | Plant | MQ | Bacteria; Gemmatimonadota; Gemmatimonadetes; Gemmatimonadales; Gemmatimonadaceae | 64.48 | 6.41 | 3359397 | 3472 |
| 3300036872_8 | Wet | Wet | Plant | MQ | Bacteria; Nitrospirota; Thermodesulfovibrionia; Thermodesulfovibrionales | 56.04 | 0.36 | 1448797 | 1708 |
| 3300036993_3 | Wet | Wet | No Plant | MQ | Bacteria; Verrucomicrobiota; Verrucomicrobiae; Chthoniobacterales; UBA10450; UBA10450 | 74.22 | 4.20 | 3077671 | 3555 |

[a]Binning was performed using the Joint Genome Institute Integrated Microbial Genomes and Microbiomes (IMG) pipeline (MetaBAT, CheckM, GTDB, GTDB-tk). Bins are accessible on IMG, under the Bin ID.

[b]Medium Quality (MQ): ≥ 50% completion and < 10% contamination.

[c]High Quality (HQ): > 90% completion and < 5% contamination.
